# Effect of Hexane and Methanol Extracts of *Piper guineense* (Schum and Thonn) Seeds against the Larger grain borer (*Prostephanus truncatus*) in Stored Cassava Chips

**DOI:** 10.1101/2024.06.17.599383

**Authors:** Emmanuel Odii, Nnaemeka Joe Okonkwo, Ibeabuchi Uko

**Affiliations:** Department of Crop Science and Horticulture, Nnamdi Azikiwe University, Awka

**Keywords:** *Prostephanus truncatus*, Hexane extract, Methanol extract, *Piper guineense*, cassava chips

## Abstract

Methanolic and hexane extract of black pepper *Piper guineense* Schum & Thonn was evaluated for its insecticidal property against the larger grain borer *Prostephanus truncatus* in stored cassava chips. The extracts were applied at various concentrations (4000, 2000, 1000, 500, 250 *µ*l/ml) with ordinary acetone as control. The experimental setup involved placing ten unsexed adults of the test insect into petri dishes treated with each of the extracts as applicable. Also, cassava chips weighing twenty grams were treated with the different concentrations of the extracts and the control before being artificially infested with *Prostephanus truncatus* and left for a period of thirty-three days. All of the treatments significantly reduced emergence holes and grain damage compared with the control.

## 1.0 Introduction

Cassava is a widely cultivated tropical root crop and the most important root crop in the tropical region (Braimah and Popoola, 2019; Amelework and Bairu, 2022). It is among the leading food crops around the world (Ray *et al*., 2019). Cassava is highly tolerant and renowned for its wide ecological adaptability, always performing relatively well where other crops may not be able to produce reasonable yield (Otekunri and Sawicka, 2019; Ikuemonisan *et al*., 2020; Berhanu, *et al*., 2023). This attribute confers on cassava a role of significant importance in ensuring food security for farming households in the tropics (Ikemonisan *et al*., 2020), in addition to providing dietary energy for nearly a billion people worldwide (FAO, 2018; Ikemonisan *et al*., 2020). Global cassava production amounted to about 278 million metric tons in 2018, with Africa contributing about 61% of the total production (FAOSTAT, 2020). Nigeria stands out as the largest cassava producer in the world, experiencing a tremendous increase in output from 7.4 million metric tonnes in 1961 to 60 million metric tonnes in 2017 (Ikuemonisan *et al*., 2020; FAOSTAT, 2022). It is one of the fastest expanding staple food crops in Nigeria, serving as a major source of dietary energy for low income consumers (Adediyan, 2015; Okoye *et al*., 2021). According to (FAOSTAT, 2020), it is projected that by the year 2025, about 62% of global cassava production will be from sub-Saharan Africa.

One of the significant challenges in cassava production is insect infestation, which extends beyond the field and continues even in storage. Insect pests gain access to the stored cassava from the standing crop in the field to various stages of grain processing and storage, resulting in seed weight loss, quality deterioration and diminished market value (Akowuah *et al*., 2018). A major destructive insect pest of dried cassava in storage is the larger grain borer, *Prostephanus truncatus*, which reduces its quantity and quality after a few months of storage (Braimah and Popoola, 2019). Despite concerted efforts by national programmes and international agencies to combat the relentless advance of *P. truncatus*; its advance in Africa has been relentless (Quellhorst, 2021). This has been a result of interstate grain trade, normal beetle flight activity and the pests’ ability to survive and breed outside storage environment. The use of chemicals for fumigation or seed treatment has become less attractive due to costs and environmental concerns. Also, challenges such as genetic resistance by insect species, pest resurgence, residual toxicity, phyto-toxicity and widespread environmental hazards have emphasized the need for alternative, organic solutions that are readily available, cost-effective and relatively non-toxic (Gilland Gard, 2014; Moshi and Matoju, 2017). These concerns have resulted in the ban of some class of pesticides, legal restrictions on some pesticides use and the desire for the development of ecological friendly pest management strategies (Mvumi and Stathers, 2014). While botanicals have shown promise in pest management, there is a need for further investigation into their efficacy, safety and standardization for practical application. The study was aimed at assessing the effect *Piper guinense* (West African black pepper) seed extracts on the weight loss and level of damage caused by *Prostephanus truncatus* in stored cassava chips.

## 2.0 Materials and Methods

### 2.1 Culture of *Prostephanus truncatus*

The insect pest culture (*Prostephanus truncatus*) was acquired from infested cassava chips purchased from Eke-Awka market and reared in the laboratory in preferred substrate of cassava chips from the same market. The transparent plastic containers used for the rearing were covered with fine mesh nets to prevent the escape or entrance of insects. The rearing containers were allowed to stand for eight weeks at room temperature to yield enough adults of the insects before they were used for the experiments.

### 2.2 Collection of Plant Materials

Dried seeds of *Piper guineense* (Schum & Thornn) were procured from Eke Awka market. This plant material was identified and authenticated at the Botany Laboratory of the Nnamdi Azikiwe University, Awka (NAUH-16). Thereafter, they were dried under room temperature for seven days and pulverized with an electric blender to obtain fine powder.

### 2.3 Soxhlet extraction from *Piper guineense* seeds

Both the methanolic and the hexane extraction were conducted using the Soxhlet extractor. In a flat bottomed flask, 250 ml of methanol or hexane was added as applicable and 40g of the milled seeds was placed in the thimble. Hot water bath was used as the source of heat. This process of extraction continued until the colour of the methanol in the extractor became clearer and the solution in the flask darker. This first packet of the milled seeds was removed and another added. The procedure continued until enough extract is gotten. Each extraction of 40g of milled pepper seeds lasted for about 4-5 hours.

### 2.4 Distillation

A water bath was used to gently heat the distillation flask containing the hexane and methanolic extracts. The water bath provided controlled and uniform heating, which is essential for safety and effectively evaporating the solvents without overheating the extracts. After the distillation process (to evaporate the solvents), a thick brownish oil with characteristic piper odour smell was obtained.

### 2.5 Preparation of Different Concentrations of the Plant Extracts

The serial dilution of the crude extract was prepared to obtain 400%, 200%, 100%, 50% and 25% concentrations thus obtaining 4000*µ*l/ml, 2000*µ*l/ml, 1000*µ*l/ml, 500*µ*l/ml & 250*µ*l/ml per 10 ml respectively with ordinary acetone as the control.

### 2.6 Residual Application of the Extracts to Determine Lethal Doses

No. 1 Whatmann filter paper measuring 9 cm in diameter was placed in each of the Petri-dishes used for the experiment. The various concentrations of the five levels of the extracts to be used were replicated three times. Control treatment with acetone was included. Subsequently, ten unsexed adults of *P. truncatus* were introduced into each of the petri dishes already treated with the hexane and methanol extracts respectively. Each of the petri dishes was covered with its lid to prevent the escape of insects. Daily mortality counts of each of the setup were taken for seven days. Both moribund and dead insects were counted to determine mortality rates.

### 2.7 Treatment of Cassava Chips

Twenty grams (20g) of cassava chips was weighed into plastic containers covered with a cap that had its center carved open and sealed with a net to allow for ventilation and prevention of entrance of insects. One milliliter each of the different concentrations of the different extracts was introduced into each of the plastic containers with the food materials accordingly. The controls were maintained with one milliliter of acetone introduced into them. Each treated plastic container was gently shaken to ensure uniform distribution of the test materials. The treatments were then left for twenty four hours to allow for proper distribution of extract and acetone to evaporate, after which the insects were introduced. Ten unsexed adults of *P. truncatus* were introduced in each of the different treatment of each extract, including the control. Before this time, the lids of the plastic vials were perforated and covered with muslin cloth held in place by a rubber band to secure the mouth of the plastic vials to ensure aeration as well as to avoid entry or exit of insects. The whole set up was then left for thirty three days. At the end of this period, records of number of emergence holes and weight loses were taken.

### 2.8 Assessment of Damage due to *Prostephanus truncatus*

Number of emergence holes in the treated cassava chips was taken at the end of the 33-day period. Similarly, weight losses were recorded to evaluate the protectant ability/efficacy of the various concentrations of the extracts. Measurements to check weight losses were carried out using an electronic sensitive weighing balance to detect even the slightest weight changes.

## 3.0 Results

### 3.1 Evaluation of mortality caused by *P. guineense* on *P. truncatus*

The effect of different concentrations and extraction solvents of *Piper guineense* on the mortality of *P. truncatus* at 24, 48, 72 hours and 7 days after treatment (DAT) is presented in Table 1. The results demonstrated that both concentration levels and extraction solvents significantly influenced mortality rates (P < 0.05) at various time points. At 24 hours post-treatment, 4000 *µ*l/ml yielded the highest mortality rate of 58.3%, significantly surpassing other concentrations, followed by 2000 *µ*l/ml at 43.3%, and 1000 *µ*l/ml at 21.7%. Both 500 *µ*l/ml and 250 *µ*l/ml showed higher mortality than the control, at 3.3%, while the control had the lowest mortality rate of 1.7%. At 48 hours, 4000 *µ*l/ml reached 100% mortality, with 2000 *µ*l/ml, 1000 *µ*l/ml, 500 *µ*l/ml, and 250 *µ*l/ml recording 51.7%, 55.0%, 6.7%, and 5.0%, respectively, while the control had the lowest mortality at 3.3.%. By 72 hours, 4000 *µ*l/ml maintained 100% mortality, while 2000 *µ*l/ml and 1000 *µ*l/ml had 58.3% and 61.7% mortality, respectively. The lowest rates persisted at 500 *µ*l/ml (10.0%), 250 *µ*l/ml (5.0%) and the control (3.3%). At 7 DAT, 4000 *µ*l/ml continued to exhibit 100% mortality, followed by 2000 *µ*l/ml at 68.3%, and 1000 *µ*l/ml at 65.0%, with 500 *µ*l/ml and 250 *µ*l/ml remaining low at 10.0% and 5.0% and the control again showing the lowest mortality at 3.3%. Among the extraction solvents, hexane consistently showed higher mortality rates than methanol, with hexane achieving 32.2%, 41.1%, 43.3%, and 45.6% at 24, 48, 72 hours, and 7 DAT, respectively, compared to methanol’s 11.7%, 32.8%, 36.1%, and 38.3%. The effect of interaction between concentration and extraction solvent was significant (P < 0.05) at all observed time intervals.

**Table 1:** Effect of concentration and extraction solvents on percentage mortality of *P. truncatus*.

| Treatment | Mortality |  |  |  |
| --- | --- | --- | --- | --- |
|  | 24hrs | 48hrs | 72hrs | 7DAT |
| **Concentration (Conc.)** |  |  |  |  |
| C1 – 4000 µl/ml | 58.3 | 100.0 | 100 | 100.0 |
| C2 – 2000 µl/ml | 43.3 | 51.7 | 58.3 | 68.3 |
| C3 – 1000 µl/ml | 21.7 | 55.0 | 61.7 | 65.0 |
| C4 – 500 µl/ml | 3.3 | 6.7 | 10.0 | 10.0 |
| C5 – 250 µl/ml | 3.3 | 5.0 | 5.0 | 5.0 |
| C6 – acetone | 1.7 | 3.3 | 3.3 | 3.3 |
| LSD | **12.40** | **13.62** | **12.08** | **12.72** |
| **Extraction Solvent (ES)** |  |  |  |  |
| Hexane | 32.2 | 41.1 | 43.3 | 45.6 |
| Methane | 11.7 | 32.8 | 36.1 | 38.3 |
| LSD (0.05) | **7.16** | **7.86** | **6.97** | **NS** |
| **Interaction** |  |  |  |  |
| Conc X ES | 17.54 | 19.25 | 17.08 | **\*\*** |

**LSD**= Least significant difference, \*\* = 0.01 level of significance

### 3.2 Evaluation of emergence holes in cassava chips treated with *P. guineense*

The effect of concentration and extraction solvent on emergence holes on cassava chips treated with *P*.*guineense* is shown in Table 2. The results demonstrated that concentration levels significantly influenced the number of emergence holes (P < 0.05). At the highest concentration of 4000 *µ*l/ml (C1), the number of emergence holes was the lowest at 0.67, indicating the most effective inhibition of *P. truncatus* emergence. As the concentration decreased, the number of emergence holes increased: 2000 *µ*l/ml (C2) resulted in 3.33 emergence holes, 1000 *µ*l/ml (C3) had 6.67 emergence holes, 500 *µ*l/ml (C4) showed 8.50 emergence holes, and 250 *µ*l/ml (C5) exhibited the highest number of emergence holes at 16.67, apart from the control. The control group (C6), which received ordinary acetone treatment, had the highest number of emergence holes at 19.00. Regarding the extraction solvents, the results indicated that hexane and methanol had a significant impact on the number of emergence holes (P < 0.05). Hexane-treated samples had fewer emergence holes (8.06) compared to methanol-treated samples (10.22). The effect of interaction between concentration and extraction solvent was significant (P < 0.05).

**Table 2:** Effect of concentration and extraction solvents on emergence holes in cassava chips treated with *P*.*guineense*.

| **Treatment** | **Emergence Holes** |
| --- | --- |
| **Concentration (Conc.)** |  |
| C1 – 4000 µl/ml | 0.67 |
| C2 – 2000 µl/ml | 3.33 |
| C3 – 1000 µl/ml | 6.67 |
| C4 – 500 µl/ml | 8.50 |
| C5 – 250 µl/ml | 16.67 |
| C6 – acetone | 19.00 |
| LSD (0.05) | **2.028** |
| **Extraction Solvent (ES)** |  |
| Hexane | 8.06 |
| Methanol | 10.22 |
| LSD (0.05) | **1.171** |
| **Interaction** |  |
| Conc. X ES | ** |

**LSD**= Least significant difference, \*\* = 0.01 level of significance

### 3.3 Evaluation of percentage weight loss in cassava chips treated with *P*.*guineense*

The effect of concentration and extraction solvents on the percentage weight loss of cassava chips treated with *P. guineense* is shown in Table 3. The results demonstrated that concentration levels significantly influenced the percentage weight loss (P < 0.05). At the highest concentration of 4000 *µ*l/ml (C1), the percentage weight loss was the lowest at 2.99%, indicating the most effective protection of the cassava chips from *P. truncatus* damage. As the concentration decreased, the percentage weight loss increased: 2000 *µ*l/ml (C2) resulted in 5.33% weight loss, 1000 *µ*l/ml (C3) had 10.50% weight loss, 500 *µ*l/ml (C4) showed 11.66% weight loss, and 250 *µ*l/ml (C5) exhibited a higher percentage weight loss at 14.04%, apart from the control. The control group (C6), which did not receive any *Piper guineense* treatment, experienced the highest percentage weight loss at 19.02%. Regarding the extraction solvents, the table showed that hexane and methanol had a significant impact on the percentage weight loss (P < 0.05). Hexane-treated samples had a higher average percentage weight loss of 13.74%, while methanol-treated samples had a lower average percentage weight loss of 7.44%. There was no significant difference (P>0.05) in the concentration and extraction solvent interaction.

**Table 3:** Effect of concentration and extraction solvents of total weight loss in cassava chips treated with *P*.*guineense*.

| **Treatment** | **% Weight Loss** |
| --- | --- |
| **Concentration (Conc.)** |  |
| C1 – 4000 µl/ml | 2.99 |
| C2 – 2000 µl/ml | 5.33 |
| C3 – 1000 µl/ml | 10.50 |
| C4 – 500 µl/ml | 11.66 |
| C5 – 250 µl/ml | 14.04 |
| C6 – acetone | 19.02 |
| LSD (0.05) | **2.141** |
| **Extraction Solvent** |  |
| Hexane | 13.74 |
| Methanol | 7.44 |
| LSD (0.05) | **3.708** |
| **Interaction (Conc. X ES)** |  |
| Conc. X ES | **NS** |

**LSD**= Least significant difference, **NS** = Not significant

## 4.0 Discussion

In the mortality assessment, there was a significant difference in the effect of concentration levels and extraction solvents at 24, 48, 72 hours and 7 days post treatment. Higher concentrations of *Piper guineense* consistently resulted in higher mortality rates, indicating a dose-dependent toxic effect on *P. truncatus*. This aligns with findings by Akhideno (2021), where increased concentrations of botanical extract led to higher insect mortality. Ileke *et al*. (2011) also recorded similar results, noting that *Piper guineense* seed extract caused 87.5% and 100% adult mortality within 24 hours of infestation at 4 days after treatment (DAT). Oben *et al*. (2015) also recorded that hexane fraction of *Piper guineense* oil induced similar mortality on *S. orizae* with 100% mortality after five days of exposure. The choice of extraction solvent also played a crucial role in the efficacy of the treatment. Hexane proved to be a more effective solvent compared to methanol, yielding higher mortality rates at all observed intervals. The significant differences observed between hexane and methanol extracts at 24 hours, 48 hours and 72 hours indicate that the solvent used can enhance or diminish the insecticidal properties of the botanical extract. The non-significant interaction between concentration and extraction solvent at 7 days post treatment suggests that while both factors independently influence the mortality of *P. truncatus*, their combined effect does not interact in a manner that significantly alters the outcome at 7 days after treatment.

The number of emergence holes in cassava chips treated with *Piper guineense* extract showed a clear dose-dependent response, with a significant difference in the effect of concentration levels and extraction solvents. At the highest concentration (C1 - 4000 *µ*l/ml), emergence holes were minimal (0.67), indicating strong suppression of insect activity. As the concentration decreased, the number of emergence holes increased significantly. The control (C6 - 0 *µ*l/ml) had the highest number of emergence holes at 19.00, highlighting the increasing damage in the absence of treatment. Furthermore, the type of extraction solvent also influenced the number of emergence holes. Cassava chips treated with hexane-extracted *Piper guineense* exhibited fewer emergence holes (8.06) compared to those treated with methanol-extracted *Piper guineense* (10.22). In a similar experiments by Ileke and Bulus (2012), *P. guineense* extracts were found to be effective against cassava lesser grain borer. Ileke (2011) also observed that fine particle size of *P. guinnense* seeds completely protected sorghum. Since most insects breathe by means of trachea which usually leads to the surface of the body called spiracle, Ileke and Bulus (2012) explained that these spiracles might have been blocked by the powders and extracts thereby leading to suffocation.

The effectiveness of *Piper guineensee* seed extract in reducing total weight loss in cassava chips infested with *Prostephanus truncatus* varied significantly with concentration and the choice of extraction. The results showed a clear concentration-dependent relationship, with higher concentrations (C1 - 4000 *µ*l/ml) resulting in minimal weight loss (2.99%), likely due to the potent bioactive compounds present in *Piper guineense* that disrupt insect feeding behaviour and metabolic processes (Isman, 2006). The control (C6 - 0*µ*l/ml) had the highest weight loss (19.02%), highlighting the susceptibility of cassava chips to *P. truncatus* without *P. guineense* (ordinary acetone) treatment. The significant differences, supported by the LSD value of 2.141, emphasized the need for higher concentrations to reduce pest damage (Adedire, 2011).

### 4.1 Conclusion

Both hexane and methanol extracts of *P. guineense* caused mortality to *P. truncatus* and demonstrated superior efficacy in reducing emergence holes and reducing weight loss in cassava chips more than the control. Although the extraction solvent did not significantly impact weight loss, hexane extracts performed slightly better.

